# RIPK3 upregulation confers robust proliferation and collateral cystine-dependence on breast cancer recurrence

**DOI:** 10.1101/679332

**Authors:** Chao-Chieh Lin, Nathaniel Mabe, Yi-Tzu Lin, Wen-Hsuan Yang, Xiaohu Tang, Lisa Hong, Tianai Sun, Tso-Pang Yao, James Alvarez, Jen-Tsan Chi

## Abstract

The molecular and genetic basis of tumor recurrence is complex and poorly understood. RIPK3 is a key effector in programmed necrotic cell death and, therefore, its expression is frequently suppressed in primary tumors. In a transcriptome profiling between primary and recurrent breast tumor cells from a murine model of breast cancer recurrence, we found that RIPK3, while absent in primary tumor cells, is dramatically re-expressed in recurrent breast tumor cells by an epigenetic mechanism. Unexpectedly, we found that RIPK3 knockdown in recurrent tumor cells reduced clonogenic growth, causing cytokinesis failure, p53 stabilization, and repressed the activities of YAP/TAZ. These data uncover a surprising role of the pro-necroptotic RIPK3 kinase in enabling productive cell cycle during tumor recurrence. Remarkably, high RIPK3 expression also rendered recurrent tumor cells exquisitely dependent on extracellular cystine and undergo programmed necrosis upon cystine deprivation. The induction of RIPK3 in recurrent tumors unravels an unexpected mechanism that paradoxically confers on tumors both growth advantage and necrotic vulnerability, providing potential strategies to eradicate recurrent tumors.

## Introduction

While significant progress has been made for the diagnosis and treatment of primary tumors, the emergence of recurrent tumors after the initial response to treatments still poses significant clinical challenges. Recurrent breast tumors are generally incurable and unresponsive to the treatments effective for primary tumors ^1^. Several factors have shown to be associated with breast tumor recurrence, including the age when primary tumor is diagnosed ^2, 3^, lymph node status, tumor size, histological grade^4, 5, 6^, the status of estrogen receptor (ER), progesterone receptor (PR) and the expression of human epidermal growth factor receptor 2 (HER2)^7, 8, 9^. However, the molecular and genetic events that lead to tumor recurrence remain largely unknown.

To study the mechanism of tumor recurrence, genetically engineered mouse (GEM) models of recurrent breast cancers have been established. Utilizing the doxycycline-inducible system, the expression of specific oncogenes can be conditionally expressed and withdrawn in the mammary gland ^10, 11, 12, 13, 14^. In the bi-transgenic mice expressing an MMTV-rtTA (MTB) and inducible Neu (homolog of HER2) oncogene (TetO-neu; TAN), mammary adenocarcinomas can be induced by the administration of doxycycline and regressed after doxycycline withdrawal ^10, 11, 12, 13, 14^. Importantly, the recurrent tumors will eventually emerge in most mice after the expression of the oncogene is turned off ^10, 11, 12, 13, 14^. This tumor recurrent model bears significant similarities to human breast cancer recurrence in several important ways: (1) Tumor recurrence occurs over a long timeframe relative to the lifespan of the mouse, similar to the timing of recurrences in human breast cancer; (2) During the latency period before recurrent tumor formation, residual tumor cells remain in the mouse, analogous to minimal residual disease in patients; (3) The formation of recurrent tumors is independent from the initial oncogene of the primary tumors, reminiscent of the finding that recurrent tumors from HER2-amplified breast cancers often lose HER2 amplification and become unresponsive to HER2 inhibition; (4) Recurrent breast cancer is often more aggressive than the initial primary tumor and resistant to therapies that were effective against the primary tumor. Breast cancer recurrence in human and GEM also share significant similarities in molecular pathways and clinical courses. For example, recurrent tumor cells of GEM typically acquired an epithelial-to-mesenchymal transition (EMT) phenotype, a hallmark of breast cancer recurrence ^13, 15^. Unfortunately, while many signaling pathways are found to be enriched in “recurrent tumor”, most of these recurrent-enriched pathways are not readily amenable to therapeutic intervention. Therefore, there are still significant need for novel therapeutic approaches that target the recurrent tumor cells.

One relative unexplored aspect of recurrent tumor is the metabolic reprogramming and potential nutrient addiction. Previously, by systematic removal of individual amino acids we demonstrated that renal cell carcinomas and triple-negative breast cancer cells (TNBC) are highly susceptible to cystine deprivation or inhibitors of cystine/glutamate antiporter (xCT) that block the cystine import ^16, 17^. Although cystine is not an essential amino acid, the imported cystine is broken down to cysteine, the limiting precursor of glutathione (GSH). GSH is a crucial antioxidant to decrease reactive oxygen species (ROS) in cells ^18^. Therefore, depletion of cystine will result in the depletion of GSH, unopposed surge of ROS, which triggers programmed necrosis ^19^. Cystine deprivation activates the Receptor Interacting Serine/Threonine Kinase 1 (RIPK1), which recruits and promotes RIPK3 autophosphorylation. The activated RIPK3, in turn, leads to the phosphorylation and polymerization of Mixed Lineage Kinase Domain Like Pseudokinase (MLKL), resulting in the membrane rupture and execution of necrosis ^20, 21^. Accordingly, as a part of death-evading strategy, RIPK3 expression is often silenced in primary tumors due to the promoter methylation ^22, 23^.

Here we report that the RIPK3 is re-expressed in recurrent tumor cells and RIPK3 activity is required for productive proliferation of recurrent tumor cells. However, this exaggerated re-expression of RIPK3 also renders the recurrent tumor cells uniquely vulnerable to the necroptotic death triggered by cystine deprivation and xCT inhibitor treatment. Thus, RIPK3-depenent proliferation of recurrent tumor cells creates the collateral vulnerability to cystine deprivation that can serve as a strategy to exterminate recurrent tumor cells therapeutically.

## Results

### Exaggerated expression of RIPK3 in the recurrent breast tumor cells

To investigate the basis of phenotypic differences between primary and recurrent tumors, tumor cells were isolated and expanded from HER2 driven murine MTB/TAN model before the oncogenic withdrawal (primary tumors) and after the recurrence (recurrent tumors)^12^. To identify differentially expressed genes important for tumor recurrence, we have previously performed microarrays to compare the differences in the transcriptome landscape between primary and recurrent tumor cells (GEO: GSE116513)^24^. This comparison validates the previously reported upregulation of Ceramide Kinase (*Cerk*) ^25^ and downregulation of Prostate Apoptosis Response 4 (*Par-4*) ^26^ in the recurrent tumor cells (Figure 1A). Also, recurrent tumor cells expressed a higher level of EMT-driving Snail Family Transcriptional Repressor 1 (*Snai1*), ^13^(Figure 1A), consistent with an enrichment of EMT by Gene Set Enrichment Analysis (GSEA) (Figure 1B). Therefore, these data confirm many distinct gene expression patterns reported between the primary and recurrent tumor cells.

**Figure 1.**
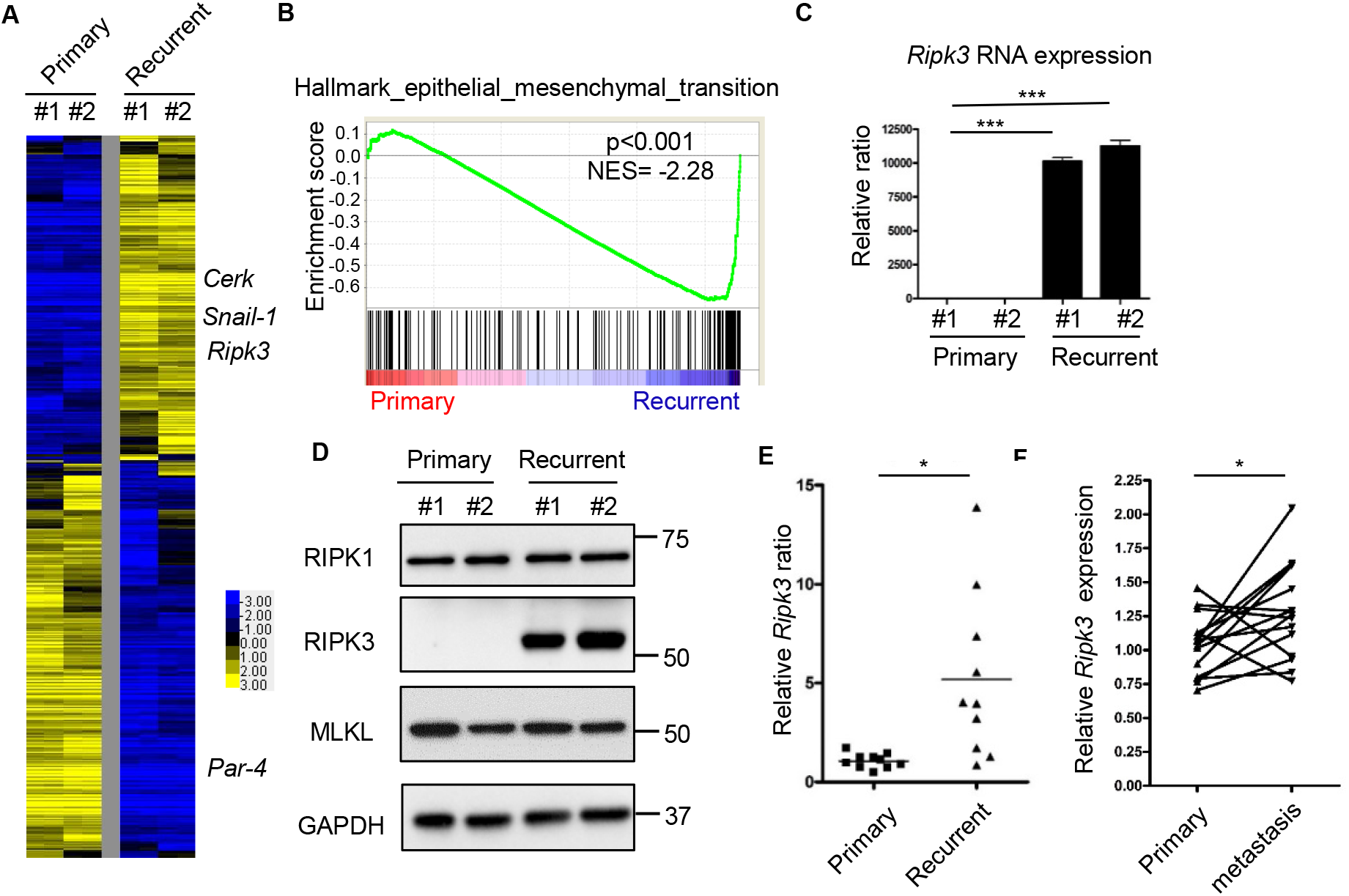
Transcriptome profiling of primary and recurrent tumor cells revealed RIPK3 upregulation in recurrent cells. (**A**) Heatmap of the transcriptional difference between two primary and two recurrent cell lines. Color scale indicates log2-fold-change. (**B**) GSEA analysis showed the enrichment of EMT geneset in the recurrent tumor cells. (**C**) *Ripk3* was highly expressed in recurrent tumor cells by RT-PCR. (**D**) Western blot showed a robust RIPK3 protein expression only in recurrent tumor cell lines. (**E**) Comparison of *Ripk3* RNA expression between 10 primary and 10 recurrent mouse tumors showed an overall increase in recurrent tumors. (**F**) Comparison of *RIPK3* expression between primary breast cancer and matching lymph node metastasis in human dataset (GSE61723). Bars show standard error of the mean. \**p* < 0.05, \*\*\**p* < 0.001, two-tailed Student’s *t*-test.

When we examined the expression of genes involved in the programmed necrosis^20^, we noted a consistent and robust over-expression of *Ripk3* in the recurrent tumors cells (Figure 1A). RT-PCR validated the dramatically increased expression of *Ripk3* mRNA in the recurrent tumor cells (Figure 1C). In addition, Western blots revealed that RIPK3 protein, while almost entirely absent in the primary tumor cells, was abundantly expressed in the recurrent tumor cells (Figure 1D). In comparison, RIPK1 and MLKL proteins, the best recognized upstream regulator and downstream target of RIPK3, respectively, were found to be expressed at similar levels in the primary and recurrent tumor cells (Figure 1D). Similar *Ripk3* mRNA over-expression is also noted in a panel of mouse recurrent breast tumors, when compared with primary breast tumors (Figure 1E). The absence of RIPK3 protein expression in primary tumor cells was previously noted and assumed to be an evolutionary strategy of tumors to escapes programmed necrosis as part of the cancer hallmarks ^22, 27^. Therefore, the absence of RIPK3 in primary tumor cells is consistent with these reports. However, the re-expression of RIPK3 in the recurrent tumor cells was unexpected.

This observation is supported by two human dataset of gene expression comparison between primary breast cancer and matching lymph node metastasis ^28, 29^. First, *RIPK3* mRNA was significantly increased by 2.08-fold in metastatic tumors when compared with primary human tumors (Supplemental Table 1)^28^. Another human dataset (GSE61723)^29^ that compared 16 pairs of primary breast cancer and matching lymph node metastasis, also showed an increase in *RIPK3* mRNA expression in 11 out of 16 pairs with an overall significant upregulation (Figure 1F). Collectively, these data indicate the upregulation of RIPK3 expression occurs in recurrent tumors in both a mouse model and two human studies.

### Epigenetic regulation of Ripk3 in the primary vs. recurrent tumor cells

To understand the basis of the *Ripk3* mRNA upregulation in the recurrent tumor cells, we investigated the epigenetic landscape of the regulatory regions of *Ripk3* gene in the primary and recurrent tumor cells by ChIP-sequencing. Consistent with the transcriptional upregulation in the recurrent tumor cells, RNA Polymerase II dramatically occupied the regulatory regions of *Ripk3* gene in the recurrent tumor cells, but not in the primary tumor cells (Figure 2A). Next, we compared the ChIP-seq data of activating epigenetic histone markers, H3K9Ac and H3K4me3, in the regulatory regions of *Ripk3*. We found that the regulatory regions of *Ripk3* gene adjacent to the RNA polymerase II binding site were highly enriched for these activating histone markers in the recurrent tumor cells, but not in the primary tumor cells (Figure 2A).

**Figure 2.**
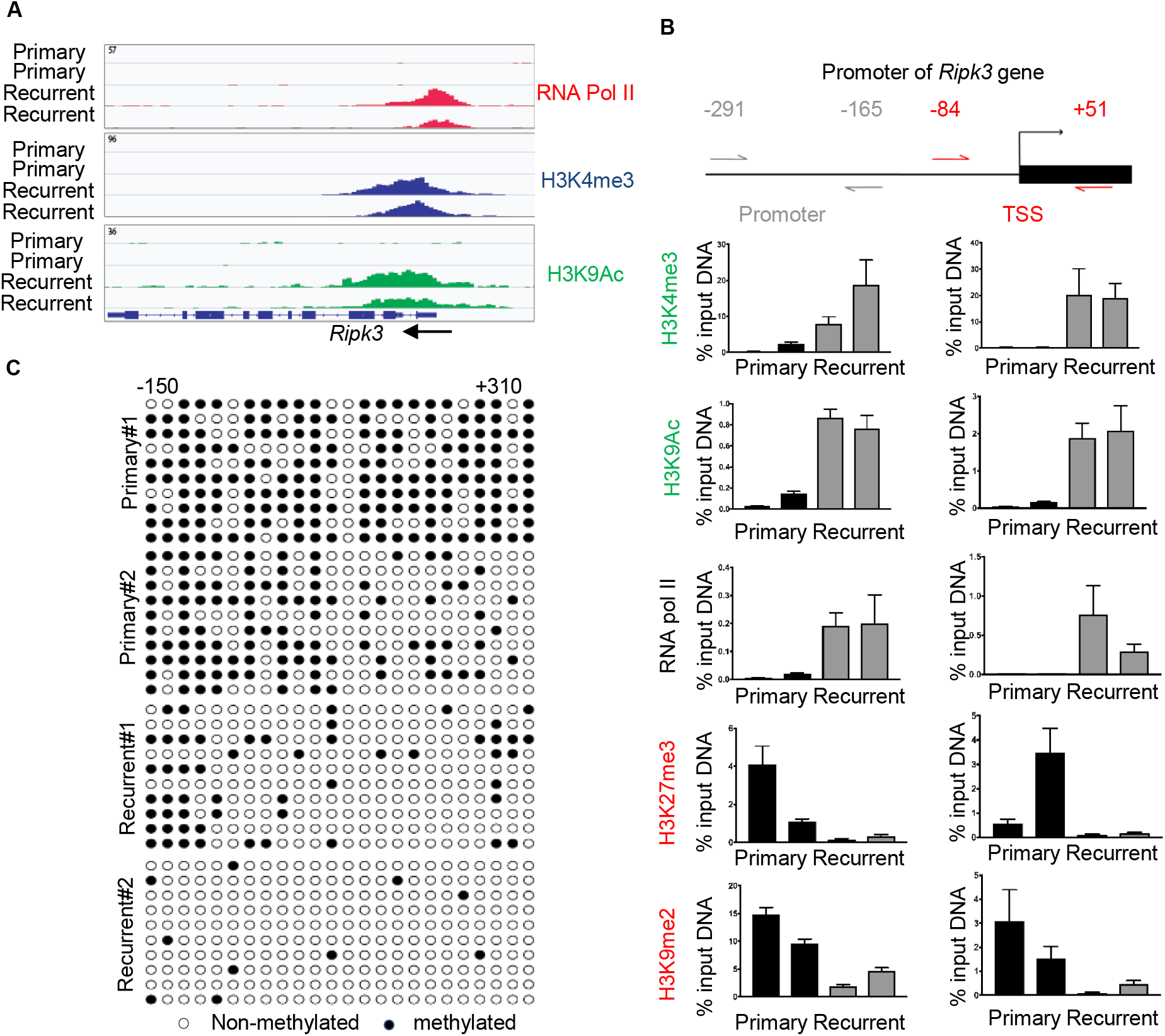
Epigenetic landscape of the regulatory regions of *Ripk3* in the primary vs. recurrent tumor cells. (**A**) ChIP-Seq data showed the occupancy of RNA Pol II, H3K4me3 and H3K9ac in regulatory regions of *Ripk3* of recurrent tumor cells (**B**) ChIP-qPCR analysis of H3K4me3, H3K9ac, RNA pol II, H3K27me3, H3K9me2 enrichment at two indicated regions in the promoter of the *Ripk3* genes in two primary and two recurrent tumor cell lines. Data are presented as the percentage of input DNA. (**C**) The cytosine methylation of CpG dinucleotides (circles) within the *Ripk3* promoter and gene body (−150 to +310) for two primary and two recurrent tumor cell lines. Bisulfite-treated DNA was transformed into bacteria and 10 replicate colonies were sequenced (rows). Open circles denote unmethylated CpG dinucleotides, while closed circles denoted methylated CpG dinucleotides.

To further determine the epigenetic alterations of *Ripk3*, we designed two sets of primers that cover the promoters (−291 to −165), transcriptional start site (TSS, −84 to +51) (Figure 2B) to measure the epigenetic changes by ChIP-PCR. We found that the promoter and TSS of *Ripk3* gene are marked by the activation markers (H3K4me3, H3K9Ac) and RNA polymerase II occupancy only in the recurrent tumor cells (Figure 2B). Reciprocally, we found that these *Ripk3* regulatory regions are marked by the silencing markers (H3K27me3 and K3K9me2) in the primary tumor cells, but not in the recurrent tumor cells (Figure 2B).

We further performed bisulfite sequencing to measure the degree of DNA methylation of the CpG island in the *Ripk3* regulatory regions (−150 to +310) (Figure 2C). We found that most of the cytosines in the *Ripk3* CpG Island are methylated in the primary tumor cells, but un-methylated in the recurrent tumor cells (Figure 2C). Together, these data indicate that epigenetic changes in the histone modification and DNA methylations are likely responsible for the silencing of *Ripk3* in the primary tumor cells and robust expression in the recurrent tumor cells.

### Ripk3 knockdown triggers mitotic defects and p53 activation

Given the unexpected robust expression of RIPK3 in the recurrent tumors, we further investigated its functional role in recurrent tumor cells. First, we performed clonogenic assay to determine whether silencing *Ripk3* in primary or recurrent tumors affects their capacity to proliferate and form colonies (Figure 3A-B, Supplemental Figure 1A-B). We found that *Ripk3* knockdown by two independent shRNAs significantly reduced colony formation in the recurrent tumor cells (Figure 3A-B), but not in the primary cells (Supplemental Figure 1A-B). These data suggest that RIPK3 is crucial for the proliferation and survival of recurrent tumor cells.

**Figure 3.**
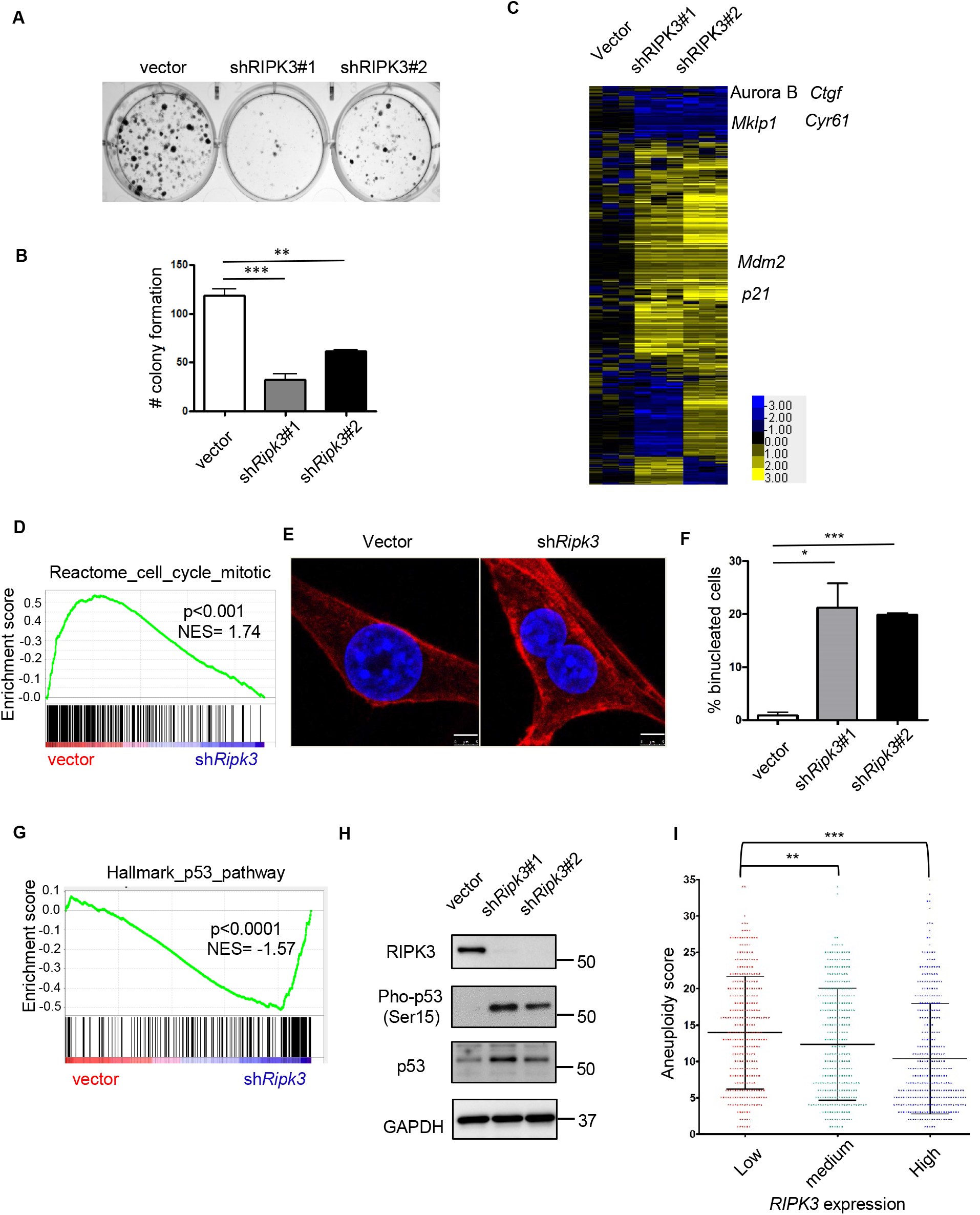
*Ripk3* knockdown triggers p53 signaling and mitotic defects. (**A**) *Ripk3* silencing decreased colony formation. Clonogenic assay was performed by plating 500 recurrent tumor cells to 6 well plates. After 10 days of incubation, cells were fixed with paraformaldehyde (4%) and stained with crystal violet. (**B**) Quantification of number of colony formation. (**C**) Heatmap of the transcriptional response to *Ripk3* silencing in recurrent cells with several affected genes indicated. (**D**) GSEA analysis showed depletion of Reactome Cell Cycle Mitosis upon *Ripk3* silencing. (**E**) *Ripk3* silencing dramatically increased binucleated cells. Recurrent cells were stained with DAPI (nucleus) and Alexa Flour 594 Phalloidin (F-actin). Scale bar, 5μm. (**F**) Quantification of binucleated cells under *Ripk3* silencing. (**G**) GSEA analysis showed that *Ripk3* silencing enriched p53 signaling pathway. (**H**) *Ripk3* silencing led to the accumulation of p53 and increased Serine 15 phosphorylation. (**I**) Lower *RIPK3* expression in human breast cancers is associated with increased amount of aneuploidy. \**p* < 0.05; **p<0.01; ***p<0.001, two-tailed Student’s *t*-test. (**B**) n=4 and (**F**) n=3 independent repeats. Bars show standard error of the mean. (**I**) n=1024.

MLKL is the downstream effector of RIPK3 in the execution of programmed necrosis ^21^. We found that *Mlkl* silencing in recurrent tumor cells recapitulated the effect of *Ripk3* knockdown and suppressed colony formation (Supplemental figure 1C-D). Consistent with these findings, treatment with an MLKL inhibitor, necrosulfonamide (NSA) ^21, 30^, also reduced colony formation of recurrent tumor cells (Supplemental figure 1E-F). These data show that the canonical necrosis-driving RIPK3-MLKL signaling axis is required for cell proliferation and clonogenic growth in recurrent breast tumor cells.

To understand the mechanisms by which *Ripk3* knockdown the reduced clonogenic capacity of recurrent tumor cells, we investigated the transcriptional responses to *Ripk3* knockdown by RNAseq (submitted to GEO: GSE124634, reviewer token: wxklcoaixtgllsp) (Figure 3C). We found downregulation of several mitotic regulators in *Ripk3* knockdown cells, including Aurora B and *Mklp1* (Figure 3C), as well as the depletion of the reactome to mitosis geneset by GSEA (Figure 3D). Therefore, we used fluorescence microscopy to investigate the potential impact of RIPK3 on mitosis. *Ripk3* knockdown dramatically increased the number of binucleated cells by ~20 folds (Figure 3E-F). These data suggest that *Ripk3* is involved in the proper execution of mitosis in the recurrent tumor cells. Binucleated/multinucleated cells generally result from cytokinesis failure ^31^. Previous study showed that cytokinesis failure can lead to the activation of tumor suppressor *p53* ^32^. By GSEA analysis, we found the enrichment of genes in *p53* signaling pathway (Figure 3G) and confirmed the upregulation of *p53* target genes, *Mdm2* and *p21*, in *Ripk3* knockdown cells (Figure 3C). Indeed, p53 protein is phosphorylated at Ser15 upon *Ripk3* silencing (Figure 3H). Phosphorylation on Ser15 led to a weak interaction between p53 and its negative regulator *Mdm2*^33^, which in terms stabilize *p53* accumulation (Figure 3H). Collectively, these data suggest that robust *Ripk3* expression in recurrent tumor cells is critical for the proper mitotic progression and cell proliferation.

Given that *Ripk3* knockdown increased binucleated cells, which cause genomic instability^34^, we speculated that *Ripk3* knockdown may lead to aneuploidy. A recent report has made the scores of aneuploidy available in a pan-cancer TCGA dataset ^35^. Therefore, we correlated the level of *RIPK3* expression with its aneuploidy score in breast cancer patients (Figure 3I). Our results indicate that low levels of *RIPK3* mRNA expression is significantly associated with higher aneuploidy in breast cancers (Figure 3I). These data in human breast tumors further support the concept that *RIPK3* is critical in preventing chromosome instability and aneuploidy in recurrent tumor cells.

### Ripk3 knockdown represses YAP/TAZ pathways

Cytokinesis failure can activate Hippo tumor suppressor pathway ^32^ and inactivate the two Hippo pathway effectors, YAP (Yes Associated Protein 1encoded by *YAP1*) and TAZ (transcriptional coactivator with PDZ-binding motif, encoded by *WWTR1*). These proteins are coactivators of TEAD family transcription factors mediating the expressions of proliferative and oncogenesis genes^36^. When Hippo is on, YAP/TAZ is inactivated by phosphorylation and exclusion from the nucleus. When Hippo is off, YAP/TAZ is localized in the nucleus and able to interact with TEAD and leads to downstream gene expression^36^. We found that *Ripk3* silencing in recurrent tumor cells led to a depletion of YAP/TAZ signature by GSEA (Figure 4A). RT-PCR confirmed that two canonical YAP/TAZ target genes: *Ctgf* and *Cyr61*, were dramatically repressed upon *Ripk3* knockdown (Figure 3C and Figure 4B). In addition, we examined how *Ripk3* silencing affects the sub-cellular localization of YAP and TAZ by nuclear/cytosol fractionation (Figure 4C). While *Ripk3* silencing slightly reduced the level of nuclear YAP, it significantly depleted nuclear TAZ (to ~18%) with a corresponding increase in the cytosolic TAZ (Figure 4C). Confocal microscopy further confirmed the reduced nuclear YAP/TAZ upon *Ripk3* silencing (Figure 4D). Thus, we speculated that the depletion of YAP/TAZ in the nucleus under *Ripk3* silencing may contribute to low efficiency of colony formation and cell proliferation. To test this hypothesis, we further over-expressed constitutively active mutants of YAP/TAZ, YAP S127A and TAZ S89A ^37, 38^ in *Ripk3* knockdown cells (Figure 4E-F). We observed a complete rescue by TAZ S89A expression under *Ripk3* knockdown whereas YAP S127A partially rescued colony formation (Figure 4E-F). These data suggest that reduced nuclear YAP/TAZ levels and activities, especially TAZ, contribute significantly to the defect of clonogenic formation induced by *Ripk3* silencing in recurrent tumor cells.

**Figure 4.**
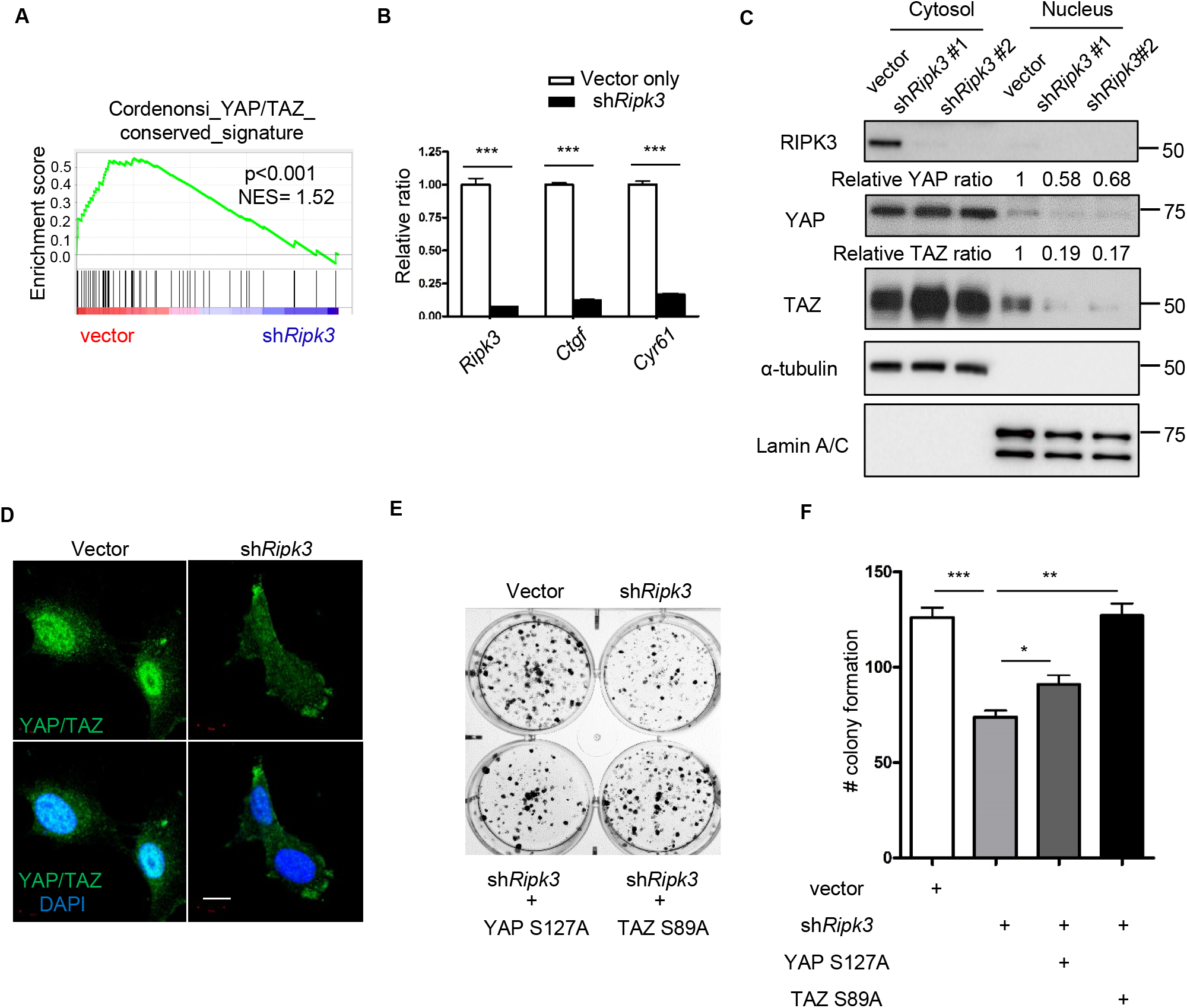
*Ripk3* knockdown abolishes YAP/TAZ-dependent cell growth. (**A**) GSEA analysis showed the depletion of YAP/TAZ transcriptional target geneset upon *Ripk3* silencing in recurrent cells. (**B**) RT-PCR validated the downregulation of *Ctgf* and *Cyr61*, two canonical YAP/TAZ target genes upon *Ripk3* knockdown. (**C**) Nuclear/cytosol fractionation showed the depletion of TAZ upon RIPK3 knockdown. α-tubulin: cytosolic marker; Lamin A/C: nuclear marker. Relative YAP/TAZ ratio was determined by normalizing YAP/TAZ intensity to Lamin A/C using ImageJ. (**D**) Confocal microscopy confirmed the depletion of YAP/TAZ upon RIPK3 knockdown. Scale bar, 10μm. (**E**) Overexpression of constitutively active YAP S127A and TAZ S89A rescued the low colony formation upon *Ripk3* knockdown as quantified in (**F**). \**p* < 0.05; **p<0.01; ***p<0.001, two-tailed Student’s *t*-test. n=3 independent repeats. Bars show standard error of the mean.

### Recurrent breast tumor cells are uniquely addicted to exogenous cystine

Recent studies have indicated that therapy-resistant and mesenchymal tumor cells, two features seen in the recurrent tumor cells, become more sensitive to cell death induced by cystine deprivation ^17, 39^. Cystine is imported into mammalian cells in exchange of the export of glutamate via the xCT transporter ^40^, which can be blocked by xCT inhibitors, such as the erastin or sulfasalazine ^41, 42, 43^. We have found that the cystine deprivation can trigger extensive cell death in renal cell carcinomas ^16^ and triple negative breast cancer cells^17^. Given the unexpected robust level of RIPK3 expression in recurrent breast cancer cells, we investigated whether recurrent breast tumor cells are particularly vulnerable to cell death triggered by cystine deprivation or erastin. We subjected two primary and two recurrent tumor cell lines to normal (200 μM cystine) or cystine-deprived (2.5 μM of cystine) media for 16 hours and determined the cell viability using crystal violet. We found that cystine deprivation eliminated most of recurrent tumor cells, but only had modest effects on primary tumor cells (Figure 5A). Under varying degrees of cystine deprivation, the recurrent tumor cells were largely eliminated under 5μM of cystine (Figure 5B). In contrast, the primary tumor cells still maintained ~50% viability even at 0.625 μM of cystine (Figure 5B). Collectively, these cell viability assays consistently showed that recurrent tumor cells are much more sensitive to cystine deprivation.

**Figure 5.**
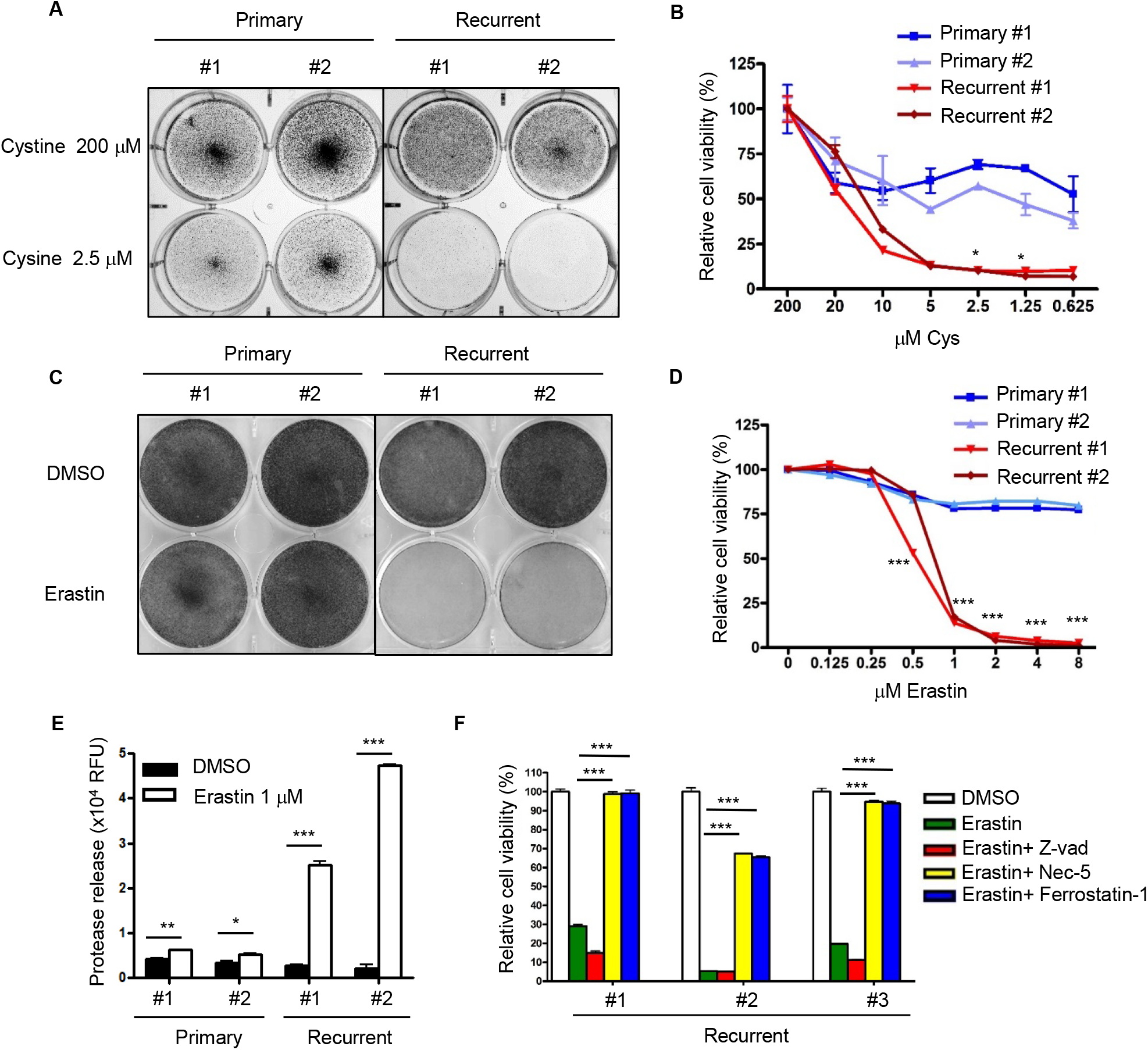
Recurrent tumor cells are more sensitive to cystine deprivation and erastin-induced death. (**A**) Recurrent tumor cells, when compared with primary tumor cells, were more sensitive to cystine deprivation. Two primary and two recurrent cell lines were incubated in full media (200 μM) or cystine-deprived media (2.5 μM) for 16 hours. The cells were then fixed with paraformaldehyde (4%) for crystal violet staining. (**B**) Recurrent tumor cells, when compared with primary tumor cells, were more sensitive to cell death under cystine deprivation. Primary and recurrent cells were incubated with decreasing level of cystine for 16 hours. The viability was then measured by ATP level using Celltiter Glo assay. (**C**) Recurrent tumor cells are more sensitive to erastin treatment than primary tumor cells. Two primary and two recurrent cell lines were incubated in erastin (1 μM) or DMSO for 18 hours. The cells were then fixed for crystal violet staining. (**D**) Primary and recurrent tumor cells were treated with increasing indicated doses of erastin for 18 hours and the viability was measured by Celltiter Glo assay. (**E**) Erastin induced more cell rupture and protease release in recurrent cells. Primary and recurrent cells were treated with 1μM of erastin for 16 hours. The media was then harvested for protease measurement. (**F**) Erastin-induced cell death was rescued by Nec-5 and Ferrostatin-1. Erastin (2 μM) were treated at the same time with DMSO, Z-vad (20μM), Nec-5 (5 μM) and Ferrostatin-1 (1 μM) in recurrent cell lines for 18 hours. The cell viability was then determined by Celltiter Glo assay. (**B,D**) *p* < 0.0001, Two-way ANOVA,, \**p* < 0.05, \*\*\**p* < 0.001. Bonferroni post hoc tests. (**E,F**) \**p* < 0.05; **p<0.01; ***p<0.001, two-tailed Student’s *t*-test. *n* = 3 independent repeats. Bars show standard error of the mean.

Alternatively, we examined whether primary and recurrent breast tumor cells have different sensitivity to erastin, a potent xCT inhibitor. Consistently, we found recurrent tumor cells, when compared with primary tumor cells, were more sensitive to erastin-induced cell death examined by crystal violet staining (Figure 5C) and CellTiter-Glo assay (Figure 5D). While recurrent tumor cells were largely eliminated between 0.5~1 μM erastin, the primary cells survived more than 8 μM of erastin with ~75% viability (Figure 5D). Such recurrent-specific erastin sensitivities are further confirmed by the higher levels of protease release in the recurrent tumor cells an indication of cell membrane breakage and death (Figure 5E).

Thus, we further analyzed the protein expression of RIPK3 and MLKL in primary and recurrent tumor cell lines after erastin treatment. We found that RIPK3 protein is only expressed in recurrent tumor cells and modestly elevated by erastin treatment (Supplemental Figure 2A). An upregulated base level of phosphorylated MLKL is noted in recurrent tumor cells when comparing to primary tumor cells (Supplemental Figure 2A). Moreover, silencing of *Ripk3* abolished the MLKL phosphorylation (Supplemental Figure 2B). Therefore, the elevated RIPK3 proteins and constitutive MLKL phosphorylation may prime the recurrent tumor cells to cell death triggered by erastin or cystine deprivation.

Erastin is considered to trigger to cell death by ferroptosis, a programmed cell death distinct from apoptosis and programmed necrosis ^42^. However, at low dose of erastin, we have previously found that necrosis pathway and RIPK3 is also required for programmed cell death ^16^. Given the elevated RIPK3 in recurrent tumor cells, we further determined the cell death mechanisms. In addition to low dose of erastin, we used different inhibitors to define the cell death mechanisms caused by erastin. We found that the apoptosis inhibitor Z-Vad did not rescue the erastin-induced death. In contrast, both ferroptosis inhibitor (ferrostatin-1) ^44^ and necrosis inhibitor (necrostatin-5) ^45^ robustly rescued the cell death, suggesting the potential role of RIPK3 in the cell death triggered by erastin (Figure 5F). While the requirement for RIPK3 may not be generally applicable to all erastin-induced cell death, RIPK3 may be particularly critical in the recurrent tumor cells with high RIPK3 expression and constitutive MLKL phosphorylation.

### Ripk3 over-expression contribute to the recurrent-specific cystine addiction

To examine whether high *Ripk3* expression in recurrent tumor cells contributes to its vulnerability to cystine deprivation, we knocked down *Ripk3* by two independent shRNAs and found a significant reduction of erastin-induced cell death with ~70-80% of viability (Figure 6A), while erastin (1 μM) eliminated the control recurrent tumor cells to less than 10% cell viability (Figure 6A). Similar results were also obtained by crystal violet staining and follow-up quantification (Figure 6B-C). Therefore, the robust *Ripk3* expression contributes to the cystine addiction phenotypes of recurrent tumor cells.

**Figure 6.**
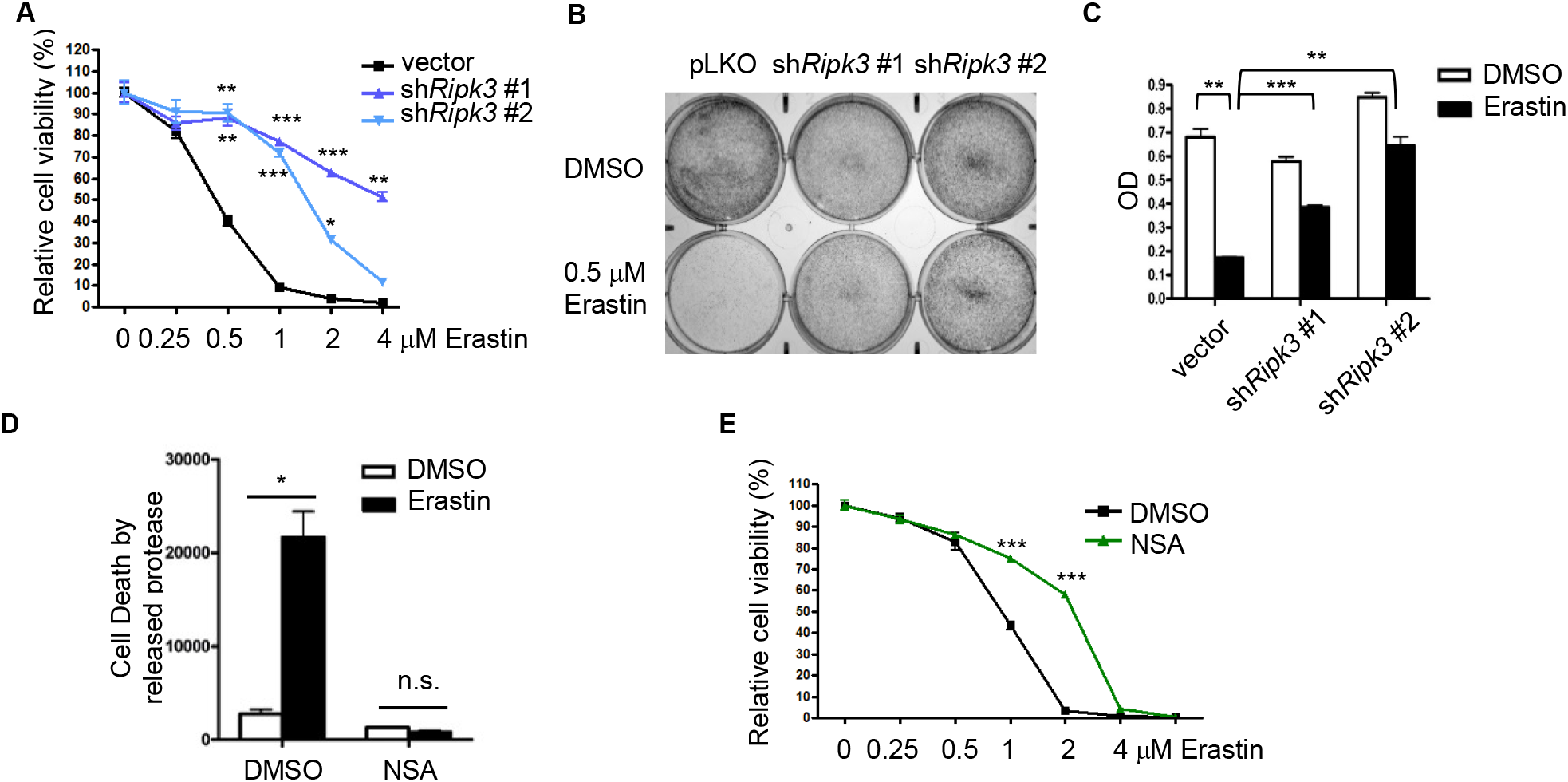
*Ripk3* over-expression contribute to the recurrent-specific cystine addiction. (**A**) *Ripk3* knockdown mitigated the erastin-induced cell death. Recurrent cells transduced with control or two *Ripk3* shRNAs were treated with increasing dose of erastin for 16 hours. Cell viability was then measured by Celltiter Glo assay. (**B-C**) Recurrent cells transduced with control or two *Ripk3* shRNAs were treated with 0.5 μM of erastin for 16 hours before assessing their viability by crystal violet staining (**b**), as quantified in(**c**). (**D-E**) MLKL phosphorylation by RIPK3 contributed to the erastin-induced cell death. Inhibiting MLKL by compound inhibitor (NSA, 5 μM) protected recurrent tumor cells from cell death under erastin treatment (0.5 μM) when measured by protease release (**D**) or Celltiter Glo assay (**E**). (**A,E**) *p* < 0.0001, Two-way ANOVA, \**p* < 0.05, \*\**p* < 0.01, \*\*\**p* < 0.001, Bonferroni post hoc tests. (**C,D**) \**p* < 0.05; **p<0.01; ***p<0.001, two-tailed Student’s *t*-test. *n* = 3 independent repeats. Bars show standard error of the mean.

Since the MLKL phosphorylation by RIPK3 is required for MLKL oligomerization and activation of necrosis, we further inhibited MLKL oligomerization by NSA ^21, 30^. We found NSA was able to rescue the erastin-induced cell death of recurrent tumor cells using either protease release assay (Figure 6D) or CellTiter-Glo assay (Figure 6E). Therefore, the exaggerated RIPK3 expression and MLKL phosphorylation, not only enable recurrent tumor cells to proliferate, but also contribute to their sensitivity to cystine deprivation and erastin treatment. These results may be limited to the recurrent tumor cells with high RIPK3 expression and constitutive MLKL phosphorylation that pose the cells to necrosis-inducing signaling triggered by cystine deprivation or erastin.

## Discussion

Global changes in epigenetic landscapes are a hallmark of cancer. While RIPK3 expression is found in most of the normal tissue, the promoter region of *RIPK3* usually becomes highly methylated in cancer cells leading to absence of *RIPK3* expression ^22, 23, 27^. Since RIPK3 determines necrosis by phosphorylating MLKL ^30^, absence of *RIPK3* expression can be considered as adaptation process for tumor cells to evade death from various necrosis-triggering signals. In this study, primary tumor cells indeed showed low *Ripk3* expression (Figure 1C-F). However, after withdrawing of oncogene, the recurrent tumor cell lines showed exaggerating amount of RIPK3 that triggers constitutive MLKL phosphorylation (Figure 1C-F and Supplemental Figure 2A); however, the activation of the RIPK3-MLKL complex does not lead to programmed necrosis (Supplemental Figure 2A). Both RIPK3 and MLKL silencing rendered recurrent tumor cells inefficient in colony formation (Figure 3A-B and Supplemental Figure 1C-D), which support the potential role of RIPK3-MLKL axis in supporting cell growth. The upregulation of RIPK3, therefore, offers selective advantages for the recurrent tumor cells.

Although the detailed mechanism remains to be discovered, our data revealed that *Ripk3* silencing increased cytokinesis failure, p53 stabilization and inactivation of YAP/TAZ signaling pathway ^32^. Thus, exaggerated expression of RIPK3 contribute to the proliferation of the recurrent cells through YAP/TAZ activities. Consistently, RIPK3 has been shown to confer survival and adaptive advantages in selected tumor settings. Knockdown of *RIPK3* in MDA-MB-231 breast cancer cells contribute to arrest in *in vivo* tumor growth ^46^. Similarly, the RIPK1/RIPK3 are highly expressed in pancreatic cancers and the in vivo deletion of these necrosis proteins delayed oncogenic progression ^47^.

Our studies also have significant therapeutic implications. Erastin can induce a non-apoptotic form of cell death named ferroptosis^42^. Although ferroptosis is considered to be regulated by glutathione peroxidase 4 and independent of RIP1/RIPK3/MLKL-mediated necrosis ^48^, our previous study has shown that low dose of erastin can induce necrosis-mediated cell death ^16^, which is supported by another study^49^. Consistent with our data, in the recurrent tumor cells with exaggerated RIPK3 expression, cell death induced by erastin can be mitigated by knockdown of RIPK3 (Figure 6). Furthermore, both Nec-5 and ferrostatin-1 rescued erastin induced cell death (Figure 5F). These data indicate that high RIPK3 expression may contribute to its sensitivities to cell death induced by cystine deprivation. While tumor recurrence is usually considered incurable, this finding suggests that the collateral vulnerability to RIPK3 mediated necrosis may hold therapeutic potential. *In vivo* cystine removal using recombinant cyst(e)inase^50^ and inhibitors of cystine importer xCT^51^ are being developed for clinical translation. Our data suggest that the recurrent tumors expressing high level of the necrosis components may be uniquely sensitive to these therapeutic approaches.

## Methods

### Cell culture

Primary and recurrent MTB/TAN tumor cells described previously^13^ were cultured in Dulbecco’s modified Eagle’s medium (DMEM; GIBCO-11995) supplemented with 10% fetal bovine serum and 1 × antibiotics (penicillin, 10,000 UI/ml and streptomycin, 10,000 UI/ml). For primary cells, 10 ng/ml EGF, 5 μg/ml insulin, 1 μg/ml hydrocortisone, 5 μg/ml prolactin, 1 μM progesterone and 2 μg/ml doxycycline were added to the media to maintain HER2/neu expression. For recurrent cells, 10 ng/ml EGF and 5 μg/ml insulin were added to the media. Both primary and recurrent cell lines were maintained in a humidified incubator at 37°C and 5% CO_2_.

### ShRNA and lentivirus infections

RIPK3 shRNA targeting mouse RIPK3 RNA were purchase from Sigma (TRCN0000022536, TRCN0000424625). MLKL shRNA targeting mouse MLKL RNA were purchase from Sigma (TRCN0000022599, TRCN000022602). Lentivirus expressing RIPK3 shRNA was generated by transfecting HEK-293T cells in 6 well plate with a 1: 0.1: 1 ratio of pMDG2: pVSVG: pLKO.1 with TransIT-LT1 transfection reagent (Mirus). After filtering through 0.45 μm of cellulose acetate membrane (VWR, 28145-481), lentivirus (250 ul) were added to a 60mm dish of recurrent cells with polybrene (8ug/ml). After 24 hours of incubation, recurrent cells were further selected with puromycin (5 μg/ml) to increase knockdown efficiency.

### Cell viability and cytotoxicity

Cell viability assay was performed by using CellTiter-Glo luminescent cell viability assay (Promega) following manufacturer’s protocol. Cytotoxicity was determined by membrane rupture and protease release using CellTox Green cytotoxicity assay (Promega) following manufacturer’s protocol.

### Western blots

Primary and recurrent tumor cell lines were harvested and washed once with ice cold PBS. The samples were then resuspended in NP-40 buffer with protease and phosphatase inhibitors and lysed by incubating in at 4°C with constant vortex for 30min, then spun down at 13000 rpm for 10 min at 4°C. Supernatant was transferred to another tube, and protein concentration was measured by BCA protein assay kit (#23225, ThermoFisher). Western blotting was performed as previously described ^52^. Nuclear and cytoplasmic extraction for YAP/TAZ was performed by following manufacturer’s protocol (#78835, ThermoFisher). Quantification of YAP/TAZ was performed by Image J software and normalized to Lamin A/C protein level. Around 20 ug of protein was loaded on 8% SDS-PAGE gels, transferred to PDVF membrane, blocked with 5% non-fat milk in 1xTBST, incubated with primary antibodies overnight at 4°C. Primary antibodies: RIPK1 (1:1000, 610458, BD biosciences); RIPK3 (1:1000, sc-374639, Santa Cruz); GAPDH (1:2000, sc-25778, Santa Cruz); Phospho-MLKL (Ser345) (1:1000, #62233, Cell signaling); MLKL (1:1000, #28640, Cell signaling); Pho-p53-S15 (1:1000, #92845, Cell signaling); Lamin A/C (1:1000, #4777T, Cell signaling); TAZ (1:1000, 560235, BD biosciences); YAP (1:1000, sc376830, Santa Cruz)

### Quantitative real-time PCR

RNA from the samples was extracted by the RNeasy Mini Kit (Qiagen) following the manufacturer’s protocol. RNA was reverse transcribed to cDNA by random hexamers and SuperScript II (Invitrogen). Quantitative real-time PCR was performed following the manufacturer’s protocol by using Power SYBR Green PCR Mix (Applied Biosystems) and StepOnePlus Real-time PCR system (Applied Biosystems). Samples were biologically triplicated for mean+/− SEM. Data were representative of three independent repeats. Mouse beta-actin (reference gene) primers: sense, 5’− GGC TGT ATT CCC CTC CAT CG −3’, antisense, 5’− CCA GTT GGT AAC AAT GCC ATG T-3’; Mouse RIPK3 primers: sense, 5’− TCT GTC AAG TTA TGG CCT ACT GG-3’, antisense, 5’-GGA ACA CGA CTC CGA ACC C-3’. Mouse CTGF primers: sense, 5’− GCC TAC CGA CTG GAA GAC AC-3’, antisense, 5’− GGA TGC ACT TTT TGC CCT TCT TA-3’. Mouse CYR61 primers: sense, 5’− CTG CGC TAA ACA ACT CAA CGA- 3’, antisense, 5’− GCA GAT CCC TTT CAG AGC GG-3’.

### ChIP-Seq and ChIP-PCR

ChIP-Seq and ChIP-PCR were performed as described previously^24^. Tumor cells were crosslinked in 1% formaldehyde (Sigma) for 10 minutes, prior to quenching with 250 mM glycine. DNA was sonicated to an average shear length of ~250-450 bp length. Lysates were precleared with protein A/G beads and immunoprecipitated with 5 μg of H3K9ac, H3K4me3, H3K27me3, H3K9me2, and RNApol2 antibodies purchased from Abcam. DNA was sequentially washed with wash buffers. DNA was eluted from washed beads and reverse cross-linked with concentrated NaCl overnight. After reverse cross-linking, proteins were digested with Proteinase K and chelated with EDTA. DNA was purified using PCR purification columns (Qiagen) according to manufacturer instructions. All qPCR reactions were carried out with SYBR green (Bio-Rad) Primers for promoter region of RIPK3: sense, 5’− CTT GGA CCC CTT AGC TCC AC-3’, antisense, 5’-GTA CCT GGC CCA AGA CAA CC-3’. Primers for TSS region of RIPK3: sense, 5’− CCC GGA CTT TGA ATG AGC GA-3’, antisense, 5’-CTC GGG TGG AAG CAG TTT CA-3’. Ct values were normalized to input DNA.

Immunoprecipitated DNA was sequenced on Illumina HiSeq 4000 sequencer with 50 bp single reads at an approximate depth of 55 million reads per sample. Sequencing reads underwent strict quality control processing with the TrimGalore package and were mapped to the mm10 genome using Bowtie aligner. Alignment files were converted to bigwig files by binning reads into 100bp segments. H3K4me3, H3K9ac and RNApol II tracks were visualized for the Ripk3 promoter by IGV desktop viewer (Broad Institute).

### Bisulfite sequencing

Bisulfite sequencing was performed using the EpiTect Bisulfite Kit (Qiagen) according to manufacturer instructions. The Ripk3 promoter region was PCR amplified using primers designed following previous study^53^. Sense, 5’− AGA GAA TTC GGA TCC TGG AGT TAA GGG GTT TAA GAG AGA T-3’, antisense, 5’-CTT CCA TGG CTC GAG CTT TAT CCC CTA CCT CAA AAA AAA C-3’. Amplified DNA was gel purified and transformed into competent bacteria. Ten independent bacterial colonies were sequenced for Ripk3, and DNA sequences were aligned with DNASTAR MegAlign software.

### Immunofluorescence microscopy

Recurrent tumor cells were washed once with PBS and fixed in 4% paraformaldehyde for 15 min, followed by permeabilization and blocking with 0.2% Triton X-100 and 2% BSA for 15 min. Primary antibodies were incubated with the cells for 1 hour. Immunofluorescence microscopy were performed using EVOS FL cell imaging system (ThermoFisher) or confocal microscope (880, Zeiss). Antibody: Alexa Fluor 594 Phalloidin (1:100, A12381, ThermoFisher); TAZ (1:100, 560235, BD biosciences); pho-histone H2AX-S139 (1:100, GTX628996, GeneTex).

### RNA-seq and GESA

TrimGalore toolkit is used to process RNA-seq data. It employs Cutadapt to trim low-quality bases and Illumina sequencing adapters from the 3’ end of the reads. Reads (>20nt) after trimming were kept for further analysis. By using the STAR RNA-seq alignment tool, reads were mapped to the GRCm38v73 version of the mouse genome and transcriptome. If reads were mapped to a single genomic location, it were kept for subsequent analysis. Quntification of read counts of genes were performed using HTSeq. Only genes that had more than 10 reads in any given library were further analyzed. DESeq2 Bioconductor package with the R statistical programming environment were applied for differential analysis to compare recurrent tumor cells with control or sh*RIPK3* silencing. The false discovery rate was calculated to control for multiple hypothesis testing. Gene set enrichment analysis was performed to identify gene ontology terms and pathways associated with altered gene expression for the comparisons between control and recurrent cells with sh*RIPK3* silencing.

### Statistical analysis

Data represent the mean +/− the standard error of the mean. P-values were determined by two ANOVA test with Bonferroni post hoc tests or a two-tailed Student’s t-test in Graphpad. Error bars represent SEM, and significance between samples is denoted as ∗P < 0.05; ∗∗P < 0.01; and ∗∗∗P < 0.001.

### Data availability

RNAseq for recurrent cells with shRIPK3 silencing has been deposited in the NCBI Genome Expression Omnibus (GEO, GSE124634). All data and reagents supporting the findings of this study are available from the authors upon reasonable request.

## Author contributions

C.C.L. and J.T.C. conceived the experiments and wrote the manuscript. C.C.L. performed the majority of the experiments. J.T.C., T.P.Y. and J.A. supervised the work. N.M., Y.T.L., W.H.Y., X.T., L.H. and T.S. collaborated in the discussion and experiments. J.T.C., T.P.Y. and J.A. provided critical feedback.

## Acknowledgments

We are grateful for technical support from the members of the Chi lab; Dr. David Corcoran for technical assistance with RNAseq. We acknowledge the financial support in part by DOD grants (W81XWH-17-1-0143, W81XWH-15-1-0486), NIH grants GM124062, the Duke Bridge Fund, Duke Cancer Institute (DCI) pilot fund.

---

**Supplemental Figure 1.**
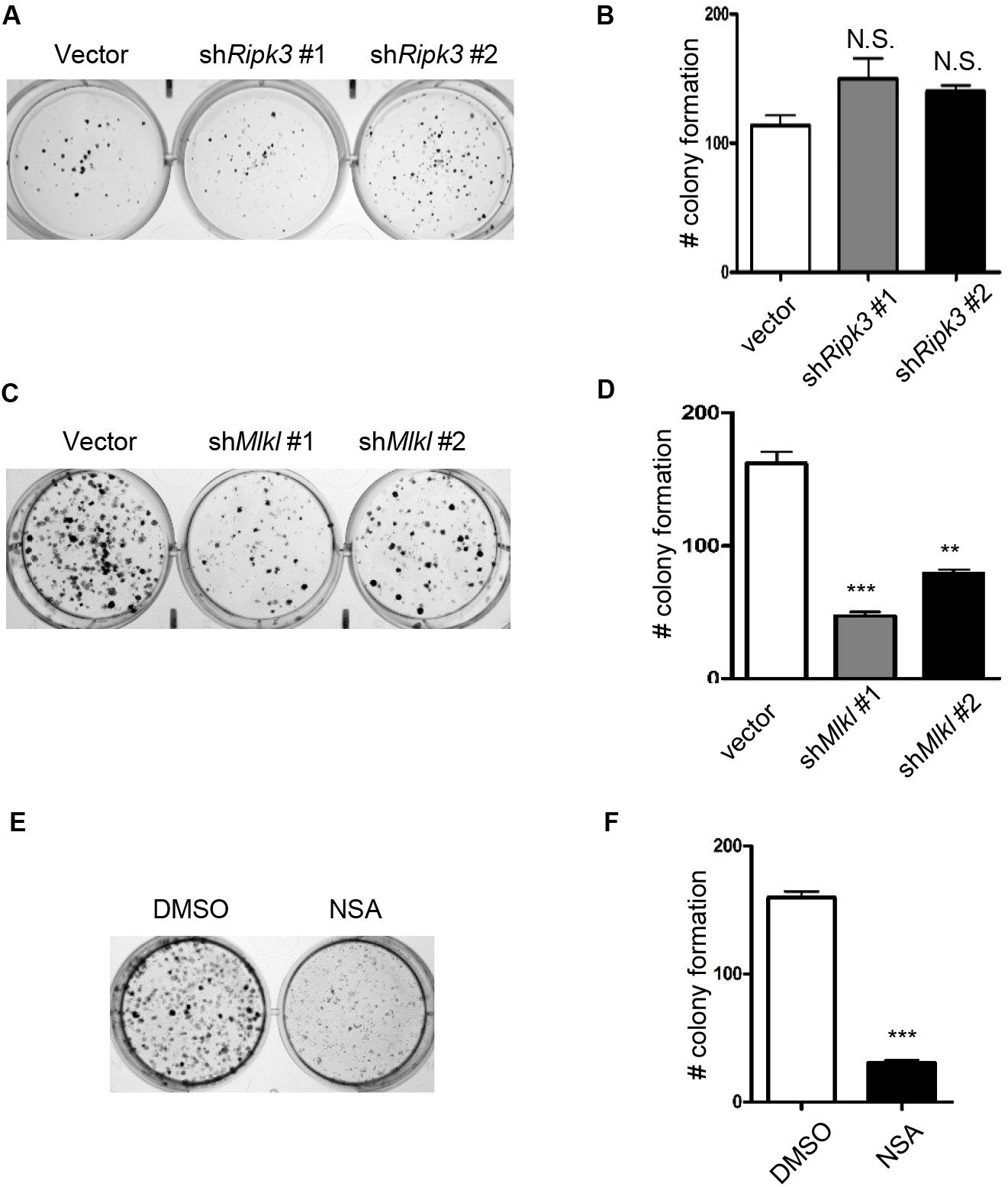
*Mlkl* inhibition by genetic or chemical means decreases colony formation in recurrent cells. (**A**) *Ripk3* knockdown did not alter colony formation in primary tumor cells. (**B**) Quantification of number of colony formation in (**A**). (**C**) *Mlkl* knockdown recapitulates the reduced clonogenic phenotype of *Ripk3* silencing in recurrent cells. (**D**) Quantification of number of colony formation in (**C**). (**E**) MLKL inhibitor (NSA, 5 μM) recapitulates the reduced clonogenic phenotype of *Ripk3* silencing. (**F**) Quantification of number of colony formation in (**E**). N.S. not significant; **p<0.01; ***p<0.001, two-tailed Student’s *t*-test. *n* = 3 independent repeats. Bars show standard error of the mean.

**Supplemental Figure 2.**
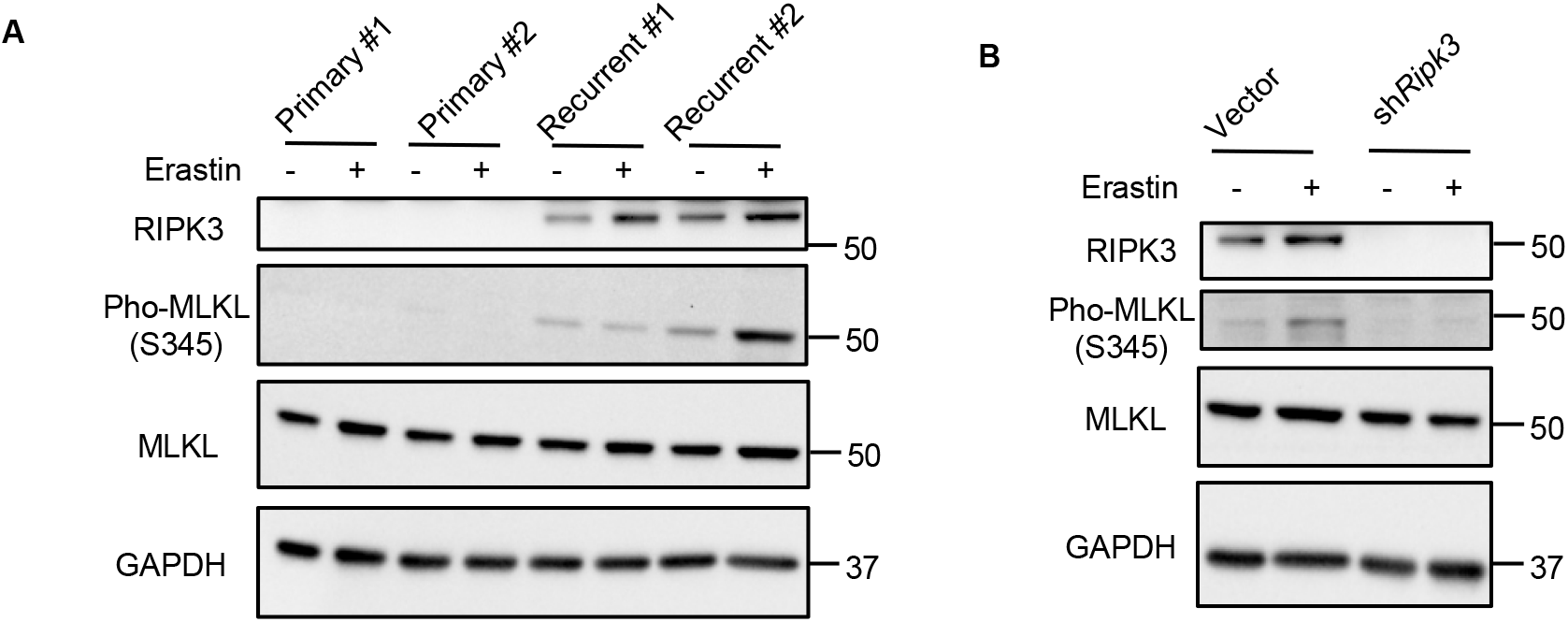
High RIPK3 expression in recurrent tumor cells is associated with the base-line phosphorylation of MLKL. (**A**) The recurrent tumor cells with high RIPK3 protein expression have higher baseline MLKL phosphorylation. Primary and recurrent cell lines were treated with 2 μM of erastin for 12 hours and lysed for Western blot with indicated antibodies. (**B**) *Ripk3* silencing abolished MLKL phosphorylation in recurrent tumor cells. Recurrent cell lines were transduced with control vector or two *Ripk3* shRNAs for 72 hours. The cells were then lysed for Western blot to measure indicated proteins.

**Supplemental Table 1.** RIPK3 expression is elevated in metastatic tumors in human dataset. RIPK3 expression in primary and metastatic tumors were evaluated by *in situ* hybridization on tissue arrays.

| Gene name | Probe Set | Genbank | fold change in metastases vs tumors | Lymph node Metastasis | Primary breast Tumor | Anova P-value |
| --- | --- | --- | --- | --- | --- | --- |
| RIPK3 | 228139_at | NM_006871 | 2.08 | 1.33 | 0.64 | 1.53E-02 |

